# WheresWalker: a pipeline for rapid mutation mapping using whole genome sequencing

**DOI:** 10.1101/2024.08.14.607950

**Authors:** McKenna Feltes, Aleksey V. Zimin, Sofia Angel, Nainika Pansari, Monica R. Hensley, Jennifer L. Anderson, Meng-Chieh Shen, Mackenzie Klemek, Yi Shen, Vighnesh S. Ginde, Hannah Kozan, Nhan V. Le, Vivian P. Truong, Meredith H. Wilson, Steven L. Salzberg, Steven A. Farber

**Author notes:** Corresponding author Correspondence to Steven A. Farber.

## Abstract

Forward genetic screening is a powerful approach to assign functions to genes and can be used to elucidate the many genes whose functions remain unknown. Chemical mutagenesis is an unbiased and efficient method for generating point mutations in the founding generation of animals in a forward genetic screening experiment. Missense and nonsense mutations induced by chemical mutagenesis can lead to the generation of partial function, gain-of-function, or null alleles that underlie compelling phenotypes, but positional cloning of the underlying causative single base pair changes can be laborious and time-consuming, especially in large polymorphic genomes. Current methods use a bioinformatic mapping-by-sequencing approach which often identifies large genomic regions which contain an intractable number of candidate genes for testing. Here, we describe WheresWalker, a modern mapping-by-sequencing algorithm that identifies a mutation-containing interval and then supports positional cloning to refine the interval which drastically reduces the number of potential candidates allowing for extremely rapid mutation identification. We validated this method using mutants from a forward genetic mutagenesis screen in zebrafish for modifiers of ApoB-lipoprotein metabolism. WheresWalker correctly maps and identifies novel zebrafish mutations in *mttp*, *apobb.1*, and *mia2* genes, as well as a previously published mutation in maize. Further, we use WheresWalker to identify a previously unappreciated ApoB-lipoprotein metabolism-modifying locus, *slc3a2a*.

## MAIN

Forward genetic screening is an established and effective technique for assigning new functions to genes, and is a particularly exciting strategy to address the ∼20% of human genes with unknown functions^1^ through screens in model organisms such as zebrafish. Chemically (e.g. ENU, EMS) induced point mutants are powerful tools for the dissection of gene function in a variety of model organisms, but relative to more-easily identifiable genome modifications (e.g., transposon-mediated insertions, CRISPR/Cas9-mediated deletions), identifying the causative single base pair substitution that underlies a particular phenotype is significantly more challenging. For the foundational chemical mutagenesis screens of the 1980-90s, mutation identification was largely achieved through a labor-intensive positional cloning approach. Specifically, mutants were outcrossed into a polymorphic background, then successively genotyped for established polymorphic markers to identify a minimal region of the genome with perfect linkage to the phenotype (recombinant mapping); that region was then amplified and sequenced^2^. Going from phenotype to mutation identification was a process that could take years. As genomic sequencing became more efficient and affordable, mapping-by-sequencing approaches emerged which utilized transcriptomic^3^, exomic^4^, or genomic^5–14^ datasets in order to bioinformatically identify mutations linked to the observed phenotype. Mapping-by-sequencing strategies range from simple genome-wide measurements of allelic frequency to more sophisticated methods that identify regions of the genome that are common in mutant animals (homozygosity mapping) or that use heterozygous alleles to predict the distance to the causative mutation^3–14^.

Both traditional positional cloning and mapping-by-sequencing approaches take advantage of the naturally occurring recombination events that shuffle the genome during germ cell generation to unlink genetic loci that originated from the same genetic origin. A centimorgan (cM) is the distance within which there is a 1% chance of recombination, and in zebrafish is ∼0.74 Mb^15^ and contains an average of 14 genes^16^. For two loci that are 1 cM apart, evaluation of 100 individuals will yield an expected 1 recombinant animal. To collect adequate whole-genome sequencing data from 100 animals as individuals or in bulk would generate very large datasets and be quite costly. Moreover, recombination frequency is non-uniform; larger numbers of animals are required to map loci positioned near centromeric regions, because those regions undergo less frequent recombination. Thus, while mapping-by-sequencing represents a massive leap forward in mutant identification, bioinformatic mapping is inherently limited by sequencing coverage. In addition, many of the existing algorithms are computationally expensive, difficult to use, unmaintained, or closed source.

We sought to develop a new pipeline that consists of 1) identification of a genomic interval linked to the phenotype using state-of-the-art alignment and SNP calling strategies and a modern and simple-to-use mapping-by-sequencing algorithm, 2) refinement of the genomic interval using traditional positional cloning, and 3) candidate gene testing using CRISPR/cas9. We call this highly efficient mapping pipeline, WheresWalker. Using whole genome sequencing data generated from pooled genomic DNA, WheresWalker compares mutants and siblings to identify the least heterozygous (most homozygous) interval in the mutant genome. To refine the interval, WheresWalker retrieves coordinates for background polymorphic markers that can be assessed in individual animals by agarose gel electrophoresis-based recombinant mapping. WheresWalker also outputs a list of candidate genes which can be tested using efficient CRISPR targeting^17^. We have applied WheresWalker to map ENU mutants with defects in apolipoprotein-B (B-lp) biosynthesis that we recently generated in a forward genetic screen in zebrafish. WheresWalker identified the correct locus for novel zebrafish alleles of *mttp*, *apobb.1*, and *mia2*. validating the strategy. To demonstrate the utility of WheresWalker beyond zebrafish, we show that the pipeline also successfully identifies a previously described mutation in maize. Finally, in a matter of weeks, we used WheresWalker to map a novel dark yolk mutant, *zion*, to *slc3a2a*, and show that this locus is linked to B-lp synthesis. Thus, we present WheresWalker as a powerful new tool for allele discovery.

## RESULTS

### Forward Genetic Screen uncovers 28 dark yolk mutants

ApoB-containing lipoproteins (B-lps) are essential for transporting lipid between tissues, but, in excess, play a causative role in a collection of metabolic disorders that impact over 1.2 billion people worldwide^18^. B-lps are synthesized in the liver and intestine in the ER lumen where microsomal triglyceride transport protein (MTP) loads lipid cargo onto ApolipoproteinB (ApoB) to form a lipid-filled particle. The ER transmembrane protein, TALI, mediates export of B-lps from the ER^19^. Like humans, Zebrafish synthesize B-lps in the liver and intestine. In addition, in larval stages, B-lps are also synthesized in the area surrounding the yolk, the yolk syncytial layer, from maternally deposited yolk lipid. As is observed in zebrafish mutants of *apobb.1* (ApoB)^20^, *mttp* (MTP)^21^, and *mia2* (TALI)^22^, disruption of B-lp synthesis results in abnormal lipid accumulation in the yolk syncytial layer which increases the opacity of the tissue. This “dark yolk” phenotype can be observed using low powered light microscopy with transmitted light.

In an effort to identify new modifiers of B-lp biology, we initiated a traditional F2 forward genetic screen looking for mutant families exhibiting the dark yolk phenotype. A founding (F0) generation of adult zebrafish males were exposed to ENU to introduce point mutations into the germ cells (Fig. 1A). A single locus hit frequency was measured in order to evaluate the mutational load of the F0 founders, using the albino locus, *slc45a2*. This locus has been used previously to evaluate mutational load in other ENU screens^4,23^ as it is an easy phenotype to screen for and is a gene of average size. In a complementation test, mutagenized males were crossed to albino *slc45a2*^b4^ females so that progeny could be evaluated for pigment defects. Of the 13,327 larvae evaluated, 17 (0.13%) exhibited the *slc45a2*^b4^ mutant phenotype (failed to complement) suggesting that 0.13% of F0 germ cells harbored a mutant copy of *slc45a2* (Fig. 1B). This level of mutagenesis is comparable to previous ENU screens in zebrafish^4,23^. Based on this analysis, we estimated that ∼800 mutagenized genomes (1/0.0013) would need to be screened in order to evaluate a deleterious mutant for each gene.

**Figure 1:**
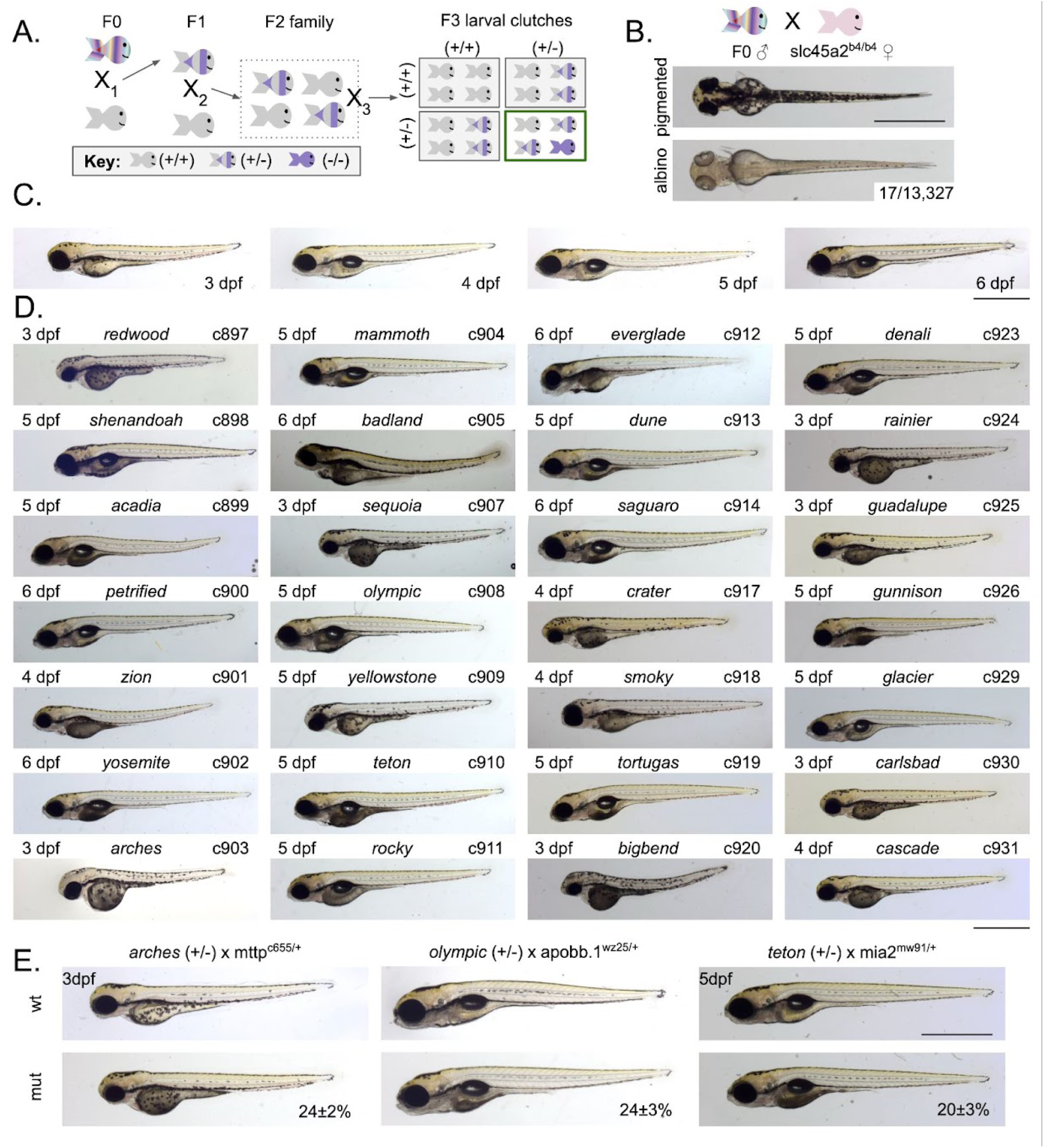
Forward genetic screen identifies 28 dark yolk mutants. A) Generation of mutant families using a standard forward genetic F3 screening scheme. B) Single locus hit rate for the *slc45a2* locus was determined by crossing male founders to *slc45a2^b4/b4^* females and screening for albinism in the offspring; representative pigmented and albino 3 days post-fertilization (dpf) larvae are shown. C) Representative images of wild-type (wt) zebrafish from 3-6 dpf. D) Representative images of identified mutants; allele ID, name, and age of animal are noted. E) Screen mutants were crossed to known dark yolk mutants so that progeny could be evaluated for dark yolk. A mutant failed to complement if dark yolk was observed at ∼25% indicating that the alleles are at the same locus. Representative images for 3 mutants that fail to complement known dark yolk loci. Phenotype frequency is reported as mean ± standard deviation. For *arches*: N = 4 clutches, n = 375 animals; For *olympic*: N = 5 clutches, n = 443 animals; For *teton*: N = 4 clutches, n = 396 animals. For all panels, scale bar represents 1 mm.

To this end, F0 founders were outcrossed to wild-type females to generate an F1 generation, each harboring 1 mutagenized genome (Fig. 1A, X_1_); individual F1 fish were outcrossed to Fus(ApoBb.1-nanoluciferase)^+/-^ to generate F2 families in a background that allows for quantification of B-lp^24^ (Fig. 1A, X_2_). Blind incrosses of F2 families were performed to generate F3 larvae which were screened for yolk phenotypes (Fig. 1A, X_3_). For each F2 family, multiple mating pairs were evaluated to increase the chance of producing progeny from a heterozygous incross which would generate homozygous progeny exhibiting recessive phenotypes (Fig. 1A, lower right quadrant); the probability of which can be calculated as 1-0.75^n^ where n is the number of mating pairs. On average, 6 mating pairs were evaluated, giving a ∼0.82 (1-0.75^6^) probability of pairing two heterozygous F2 fish in at least one of the blind intercrosses. This probability (1-0.75^n^) was used to calculate the fraction of the genome screened for each of the 1,023 families screened and summed to equal 814 genomes which represents 1.05X genomic coverage. Twenty-eight dark yolk mutants were identified (Fig. 1D), of these, 27 produced the phenotype in mendelian ratios and could be recovered in subsequent generations (Extended Data Fig. 1A).

Alleles of *mttp*^21^, *apobb.1*^20^, *mia2*^22^, *dgat2* and *pla2g12b*^25^ have already been reported to produce the dark yolk phenotype. In order to determine if these loci were represented in our mutant collection, each novel mutant was crossed to the known dark yolk alleles to generate progeny for evaluation. Using this complementation approach, 4 *mttp* alleles (*arches*, *bigbend*, *guadalupe*, *carlsbad*), 2 *apobb.1* alleles (*mammoth*, *olympic*), 2 *mia2* alleles (*teton*, *dune*), and 2 *dgat2* alleles were identified (Fig. 1E, Extended Data Fig. 1B-E); *pla2g12b* was not identified in this screen suggesting saturation was not reached. Three of these known dark yolk mutants (*mttp*(*arches*), *apobb.1*(*olympic*), *mia2*(*teton*)) were selected to be evaluated by sequencing for development of the mapping pipeline.

### SNP Index is a measure of homozygosity for bulk segregant analysis

For recessive mutations, the causative locus and surrounding genomic region will be homozygous in mutant animals, while regions outside of the locus will be more heterozygous as zebrafish are highly polymorphic^26^. To leverage this principle we designed an algorithm that utilizes whole genome sequencing (WGS) data from mutant and wild type pooled genomic DNA to identify regions of the genome that are more homozygous in mutant animals (Fig. 2). To generate WGS datasets, a bulk segregant analysis approach was undertaken. Heterozygous adults were incrossed to generate clutches containing both phenotypically wild type (+/+ and +/-) and phenotypically mutant (-/-) larvae which were sorted by phenotype and pooled before being prepared for sequencing. WGS data was aligned and evaluated for points of variance using POLCA^27^, a faster and more accurate genome polishing tool (relative to Pilon) that generates a report on genome variance in the form of variant call format (VCF) files (Fig. 2A). To identify regions of homozygosity in the mutant genome, the algorithm uses 10 Kbp sliding windows to quantify the mean degree of heterozygosity for mutant *H_m_(C)* and wild type *H_w_(C)* datasets by counting the number of heterozygous points in each window, where *C* is the coordinate of the window center (Fig. 2B). Relative homozygosity, or SNP index is calculated as follows: *SNP_index_(C)=*(*H_w_(C)-H_m_(C)*)/(2-*H_m_(C)).* In regions where mutants are less heterozygous (more homozygous) than non-mutant siblings, *H_w_(C)* will be much larger than *H_m_(C)*, causing the *SNP_index_(C)* to be high and positive. To smooth out noise we apply a moving average *MA* filter to *SNP_index_*with the window width of 1 cM. The genome then is scanned for the highest value of *MA(SNP_index_(C))* and an interval around the maximum, bounded on each side by *MA(SNP_index_(C))=*0 is selected for further analysis (Fig. 2C). SNPs are retrieved from the interval and selected if the SNP segregates appropriately with the mutant phenotype (see Methods). A modified version of ANNOVAR^28^ software is used to annotate all mutations in the window and all nonsynonymous mutations are outputted. The SIFT score^29^ is added for previously annotated zebrafish polymorphisms so that novel SNPs can be prioritized (Fig. 2D). We call this pipeline “WheresWalker”, and it is publicly available on github at https://github.com/alekseyzimin/WheresWalker.

**Figure 2:**
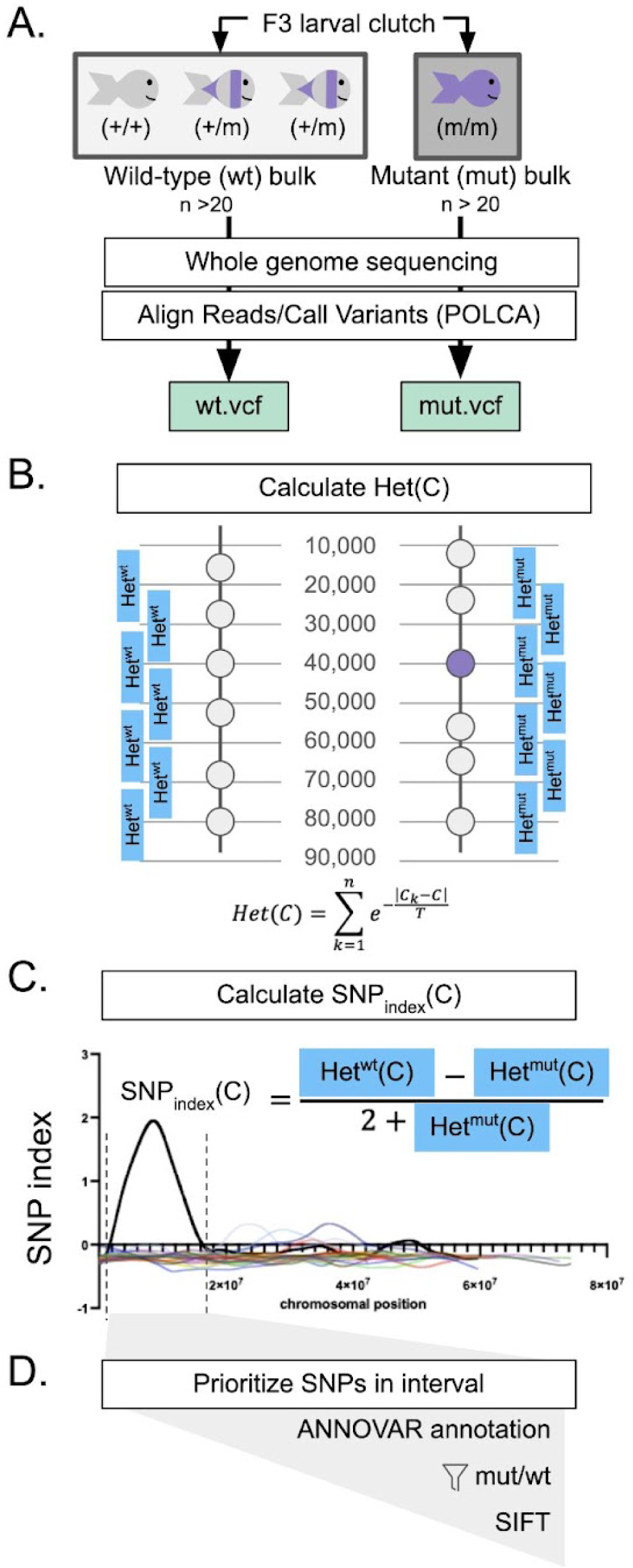
WheresWalker pipeline utilizes WGS data to identify segregating SNPs and indels. A) Bulk Segregant Analysis: animals are sorted by phenotype to generate wild-type (wt) and mutant (mut) bulk genomic DNA for whole-genome sequencing. Sequencing data is aligned and evaluated for variance using POLCA which outputs VCF files for wt and mut samples. B) Heterozygosity is calculated in a sliding window across wt and mut genomes, where C is the coordinate at the center of each window. Gray dots represent points of heterozygosity, purple points represent the causative mutation. C) wt and mut heterozygosity is compared to identify regions in mut that are more homozygous by calculating the SNP index and an interval is selected; dashed lines indicate interval bounds. D) SNPs are annotated and selected as candidates if they segregate appropriately. The SIFT score is added for selected SNPs.

### Mapping pipeline identifies correct chromosome for 3 known dark yolk loci

In order to test WheresWalker, WGS data was collected from wild type and mutant genomic DNA pools for three of the novel dark yolk alleles identified in our forward genetic screen: *mttp*(*arches)*, *apobb.1*(*olympic)*, and *mia2*(*teton)*. For each set, VCF files generated by POLCA were submitted to WheresWalker. For each of the 3 mutants, visual assessment of the SNP index for every chromosome indicated a single chromosome with elevated SNP index (Fig. 3A-B). For *arches*, a 11.40 Mb interval on chromosome 1 was selected which contained the causative locus, *mttp*. A 36.86 Mb interval on chromosome 20 containing *apobb.1* was selected for *olympic* (Fig. 3B) and a 17.56 Mb interval on chromosome 17 containing *mia2* was selected for *teton* (Fig. 3C). In all cases, the pipeline was sensitive enough to select the correct chromosome and the genomic interval of the causative locus even without an outcross.

**Figure 3:**
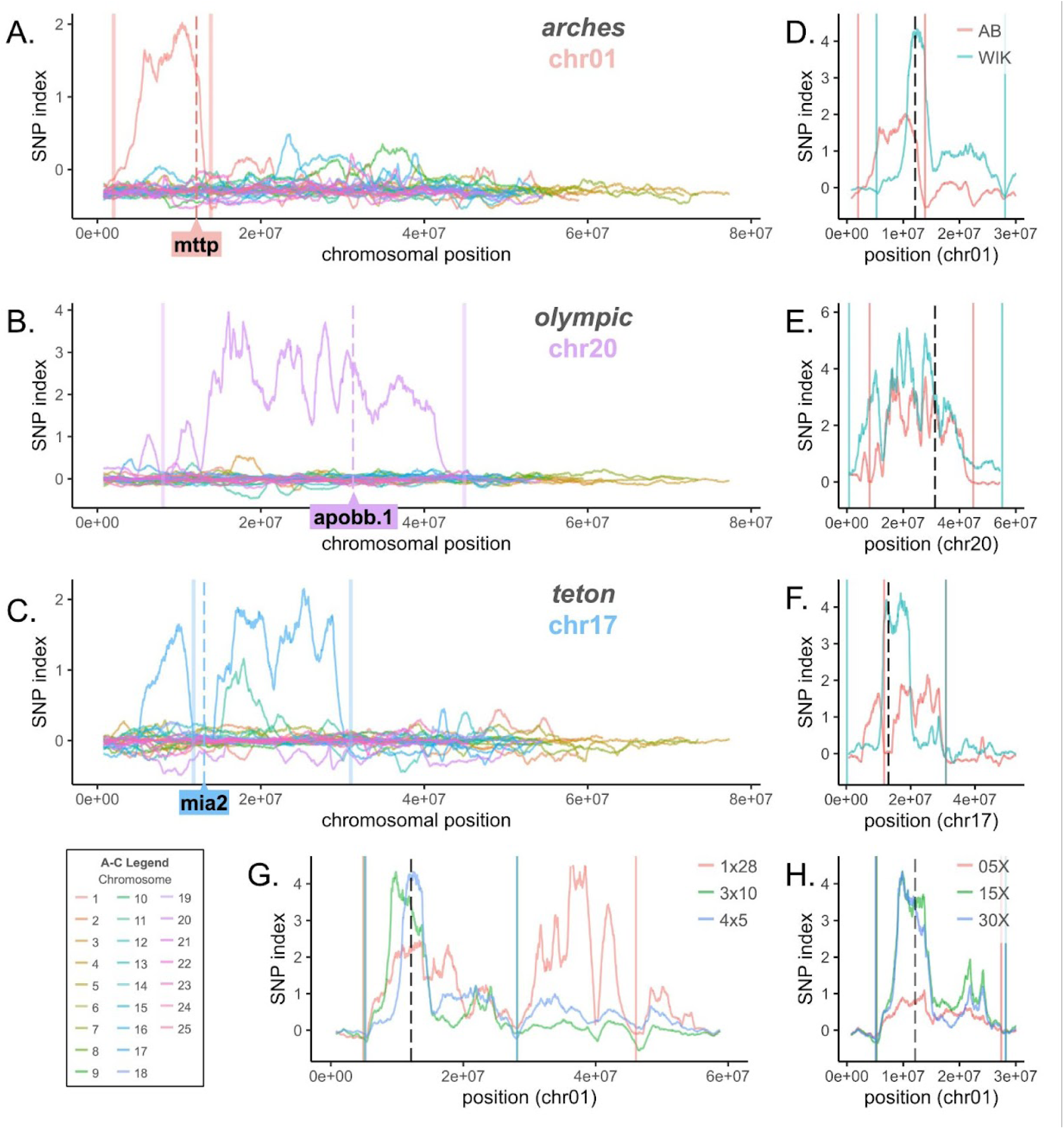
WheresWalker identifies the correct chromosomal region for three dark yolk loci. Profile of SNP index across all chromosomes for *arches* (A) *olympic* (B) and *teton* (C) mutants. Solid lines indicated left and right bounds of the interval selected by WheresWalker, dashed lines indicate the position of the causative locus. D-F) regional SNP index profile for bulks generated in an in-crossed (AB) or outcrossed (WIK) background for *arches* (D), *olympic* (E), and *teton* (F) mutants. G) Regional SNP index for *arches* bulks generated from different combinations of clutches: 1x28, 3x10, or 4x5 (clutch x animals/clutch). H) Regional SNP index profile for the *arches* WIK, 3x10 dataset with sequencing coverage simulated at 05X, 15X, or 30X. For D-H Solid lines indicate left and right bounds of the interval selected by WheresWalker, dashed lines indicate the position of the causative locus.

Conventional mapping relies on an outcross to a different wild-type strain to introduce microsatellite markers for recombinant mapping. We hypothesized that the introduction of polymorphisms in this way would also improve bioinformatic interval picking. To test this, genomic DNA pools were generated for *mttp*(*arches)*, *apobb.1*(*olympic)* and *mia2*(*teton)* mutants and siblings in the WIK background and processed using WheresWalker. As in the original datasets, the interval selected contained the causative locus for each mutant (Table 1, Extended Data Fig. 2A, F, I). Relative to intervals selected in the AB background, intervals in the WIK background were wider, however, the causative locus was more likely to be close to the center of a major SNP index peak and the mean SNP index value in the 2 cM surrounding the mutation was higher (Table 1, Fig. 3D-F).

**Table 1:**
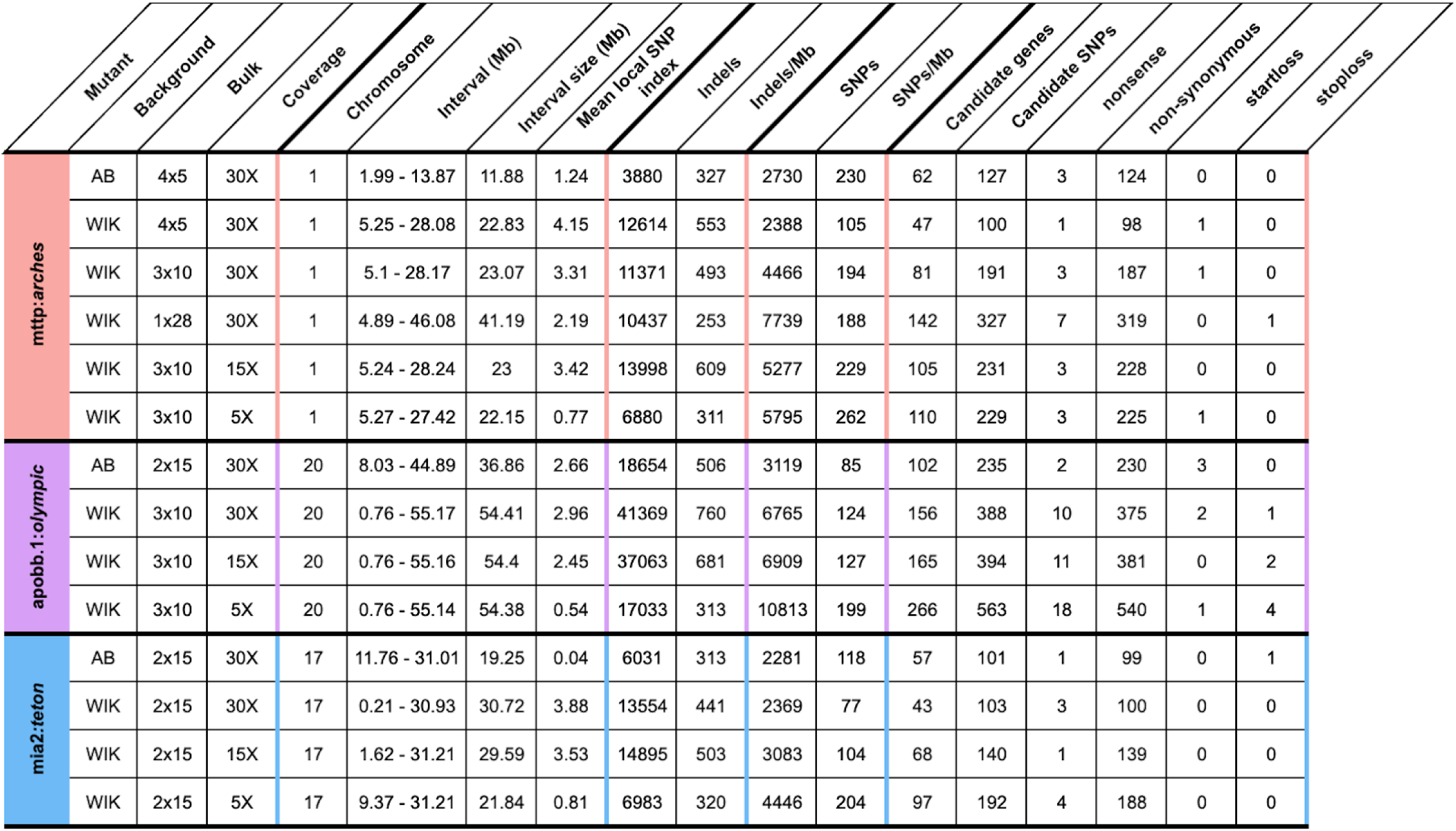
Summary of WGS datasets.

We predicted that increasing the genetic diversity represented in the bulk samples by increasing the number of clutches represented in each bulk would further improve peak quality. To test this, bulk samples from *mttp*(*arches)* were generated which represented 1 clutch (n=28, 1x28) or 3 clutches (n=30, 3x10) in the WIK background and compared them to the original WIK bulk which was made up of fewer animals but represented 4 clutches (n=20, 4x5). For each set, an interval containing *mttp* on chromosome 1 was selected (Table 1, Extended Data Fig. 2A-C), but the selected interval was almost 2X larger when only one clutch was represented. Generating bulks representing 3+ clutches decreased the interval size and increased the mean SNP index around the mutation (Fig. 3G, Table 1). To test genomic coverage requirements, existing datasets were trimmed to represent 5X and 15X coverage for the *arches* WIK 3x10 dataset. Increasing coverage above 5X did not significantly impact the size and location of the interval selected but substantially increased the SNP index in the interval and near the mutation (Table 1, Fig. 3H, Extended Data Fig. 2D, E). Increasing coverage from 15X to 30X did not change the qualitative appearance of the interval (Fig. 3H), but did reduce the number of segregating SNPs (Table 1). A similar effect was observed at *apobb.1* and *mia2* loci (Table 1, Extended Data Fig. 2F-M). The number of candidate genes in each interval was determined by selecting only genes with exonic missense or nonsense mutations (Table 1). On average, increasing coverage from 5X to 15X reduced the number of candidate genes by 24±17% and increasing coverage from 15X to 30X reduced the number of candidate genes by an additional 22±16%.

To further test the utility of WheresWalker, we applied the tool to a publicly available WGS dataset for a mutation in maize (Zea mays) called *very narrow sheath* (*vns*) which has been identified as an allele of the *defective kernel 1* (*dek1*) gene^30^. VCF files were generated from WGS data for mutant and wild type pools which were inputted to WheresWalker. Even with a small number of individuals represented in the dataset (9 for each phenotype) and moderate sequencing coverage (17X), WheresWalker was able to identify a 75.71 Mb interval on chromosome 1 containing *dek1* (Extended Data Fig. 3). The *dek1* locus was ∼ 3 Mb from the highest SNP index peak. These data demonstrate the utility of WheresWalker beyond zebrafish.

### Recombinant mapping narrows region of interest to identify a nonsense allele of *mttp*

With 30X genomic coverage, WheresWalker picks an interval of ∼10-50 Mb which represents 0.5-3% of the zebrafish genome. This is a substantial reduction in the total amount of genomic space to search for the causative SNP but is still quite large and contains hundreds of candidate genes (Table 1). In addition to SNPs, polymorphic insertions and deletions (indels) are also detected by WGS. Within the intervals selected at each loci, segregating indels were identified at 253-760/Mb (Table 1). We hypothesized that these indels could be assessed for linkage to the phenotype in order to further reduce the genomic space in consideration. A module was added to WheresWalker to extract the position and size of indels in the interval. To test this module, PCR primers were designed around indels on either side of the *mttp*(*arches)* interval and are referred to as markers “A_a_” and “B_a_”. *mttp^arches^*^/+^; A_a_^+/-^; B_a_^+/-^ parental fish were incrossed to generate *mttp^arches^* mutant progeny which were collected and genotyped for markers A_a_ and B_a_ (Fig. 4A, Extended Data Fig. 4A). Most of the 35 mutant progeny were homozygous mutant at both markers (A_a_^-/-^; B_a_^-/-^), but 12/35 for marker A_a_ and 1/35 for marker B_a_ were heterozygous indicating a recombination event occurred, unlinking the marker from the phenotype. Because recombination frequency is a function of the linear distance between two genetic loci, these results indicated that marker A_a_ was farther from the mutation than marker B_a_. Further, in the sole recombinant for marker B_a_, animal 5, recombination was also observed at marker A_a_ suggesting both makers were on the left side of the mutation. These data exclude from consideration the region of 2.46 Mb (original interval bound) to 10.71 Mb (marker B_a_) as the location of the causative locus. The remaining 3.16 Mb region (10.71-13.87 Mb) contains 52 candidate SNPs in 24 unique genes, including a single base pair change from G>T at position 12109076 leading to a nonsense mutation in *mttp*: ENSDARG00000008637: ENSDART00000015251: exon17: c.C2475A: p.C825X, which was confirmed by Sanger Sequencing (Extended Data Fig. 4B). Further, an estimation of the distance to the causative locus, calculated using the recombination frequency, predicted markers A_a_ and B_a_ to be 25 and 2 Mb away (lines, Fig. 4B). *Mttp* was within the bounds of both markers and was just 1.4 Mb from marker B_a_.

**Figure 4:**
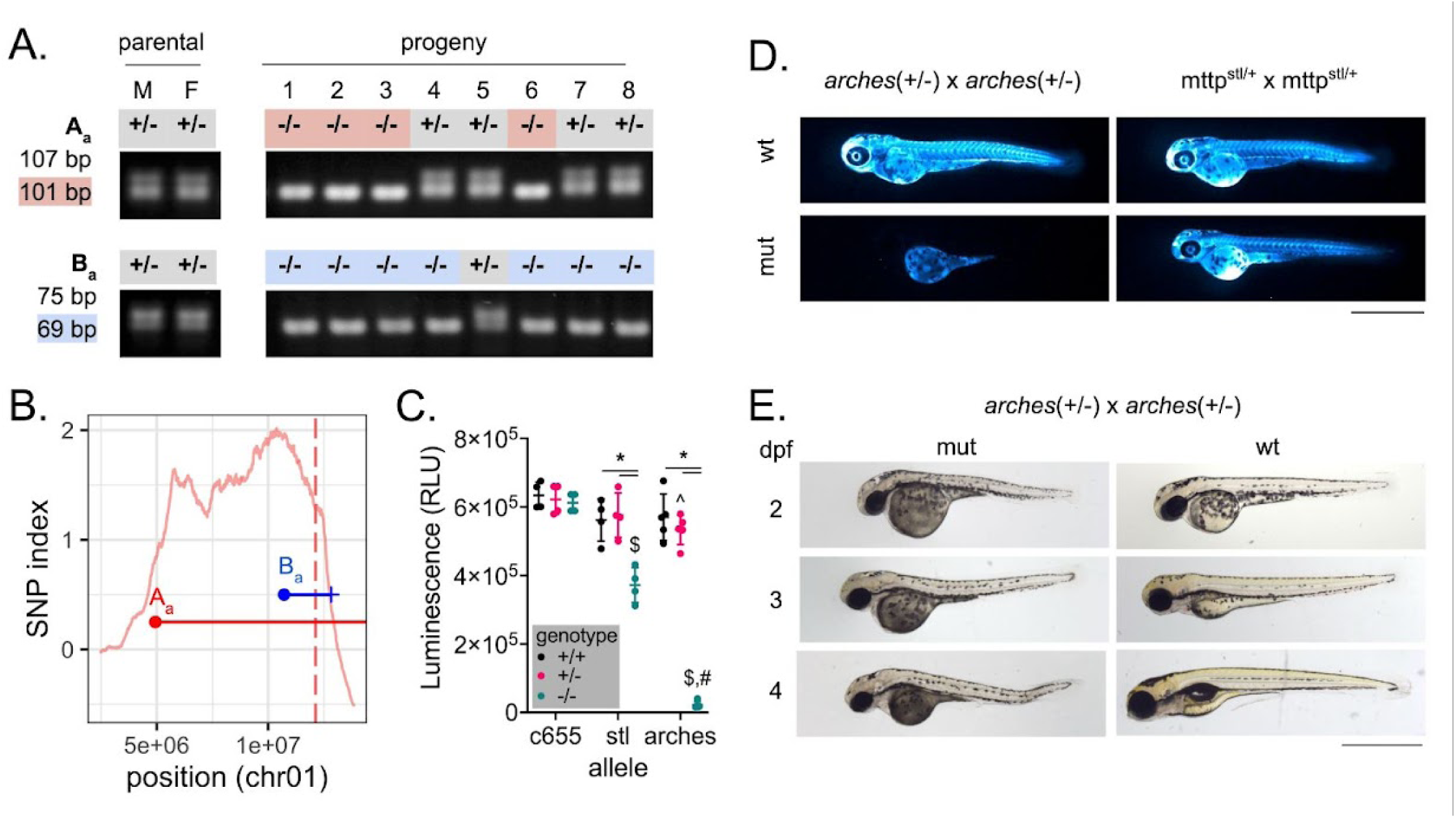
Recombinant mapping narrows region of interest to identify a hypomorphic allele of *mttp*. A) markers A_a_ and B_a_ were outputted by WheresWalker and used to genotype *arches* mutants in order to identify recombinants. M and F denote male and female parents, respectively. PCR product sizes for wild-type and mutant (highlighted) products are as indicated. B) Points representing A_a_ (red) and B_a_ (blue) marker locations and horizontal lines representing the estimated distance to the causative mutation are overlaid on the SNP index for the *arches* interval (chromosome 1: 2.46-13.86 Mb). The vertical dashed line indicates the location of *mttp*, the causative locus. C) quantification of ApoBb.1-nanoluciferase levels at 3 dpf. Mean +/- standard deviation, N = 4-5 clutches, n = 2-8 animals/datapoint. P<0.05 by two-way ANOVA with Geisser-Greenhouse correction and Tukey’s multiple comparisons test. * vs. respective wild type, ^ vs c655(+/-), $ vs c655(-/-), # vs stl(-/-). D) Images of whole-animal ApoBb.1-nanoluciferase distribution in *arches* and *stl* mutants at 3dpf. E) Brightfield images of mutant and wild-type animals from 2-4 dpf. For D and E, scale bar represents 1 mm.

The *mttp arches* mutation introduces a premature stop codon in exon 17 leading to a 59 bp truncation. A similar truncation, observed in a human patient with abetalipoproteinemia, was shown to disrupt the binding of the *mttp* protein product, MTP, with PDI which is essential for function^31^. We therefore predicted that the *arches* mutants would have a severe phenotype and sought to compare them to previously studied zebrafish *mttp* alleles *c655* and *stl* which have deficiencies in triglyceride and triglyceride/phospholipid transfer to ApoB, respectively^21^. ApoB quantity was measured using the LipoGlo reporter system^24^ in *mttp^arches^*, *mttp*^c655^, and *mttp*^stll^ mutants (Fig. 4C). At 3 dpf *mttp*^c655^ have wild-type levels of ApoB while *mttp*^stl^ levels are reduced by ∼50%. In contrast, ApoB was hardly detectable in *mttp^arches^* embryos suggesting very few B-lp particles were produced. This finding was further confirmed when fixed whole embryos were assessed for ApoB localization: observable ApoB signal was restricted to the yolk syncytial layer, the B-lp synthetic tissue during embryonic stages^24,32–34^ (Fig. 4D). While *mttp*^stl^ and *mttp*^c655^ survive to adulthood^21^, *mttp^arches^* fish exhibit yolk retention and develop tissue necrosis early in development (Fig. 4E) and do not survive past larval stages further illustrating the severity of the novel *arches* allele.

### *zion* maps to *slc3a2a*, a novel dark yolk locus

To further test WheresWalker, we selected one of the novel dark yolk mutants, *zion*, to be mapped. 30X WGS datasets were collected for wild type and mutant bulks which were inputted to POLCA^27^ to generate VCF files that were submitted to WheresWalker. The pipeline selected a 30.4 Mb interval on chromosome 7 (Fig. 5A, Table 2) which contained 150 exonic mutations in 80 genes. Indels, from the WheresWalker output, were selected and 5 markers (A_z_-E_z_) were optimized for recombinant mapping. A total of 121 dark yolk larvae from an incross of a single parental pair of *zion*^+/-^ adults were collected and genotyped for each marker (Extended Data Fig. 5A-B). Mapping reduced the region of interest to ∼7 Mb between markers D_zion_ and E_zion_ (19.03-26.12 Mb) (Fig. 5B); this region contained 42 exonic mutations in 25 genes. The recombination frequency was used to predict the distance to the causative locus from all markers, which averaged to 20.23 +/- 0.96 Mb (Extended Data Fig. 5C). This ∼2 Mb region contained mutations in 8 genes including 12 nonsynonymous and 1 nonsense SNP. The single nonsense mutation in *slc3a2a* (ENSDARG00000036427: ENSDART00000052917: exon7: c.C1012T: p.Q338X) was prioritized as the top candidate (Extended Data Fig. 6A). CRISPR guides targeting *slc3a2a* were injected into 1-cell zebrafish embryos with Cas9 to induce editing. Guides targeting the ohnolog of *slc3a2a*, *slc3a2b*, were also tested. At 4 dpf, *slc3a2a* injected larvae phenocopied the *zion* dark yolk phenotype, whereas no dark yolks were observed in *slc3a2b* crispants (Fig. 5C). Editing at *slc3a2a* and *slc3a2b* loci was confirmed by PCR (Extended Data Fig. 6B,C).

**Figure 5:**
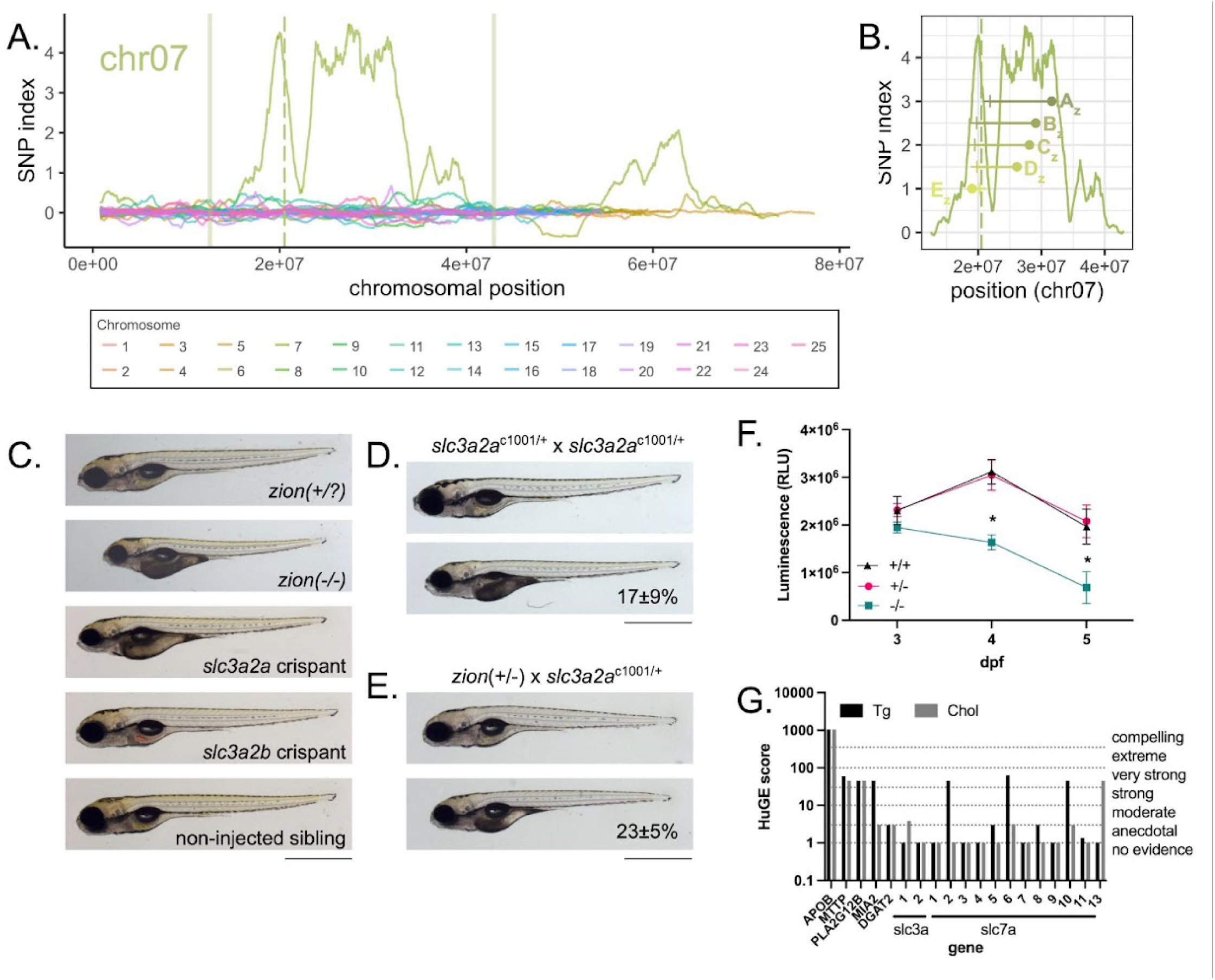
*zion* maps to *slc3a2a*. A) Elevated SNP index is observed on chromosome 7 in *zion* mutants; WheresWalker selected an interval from 19.03-26.02 Mb which was further analyzed. Solid vertical lines indicate interval bounds on chromosome 7. The vertical dashed line indicates the position of *slc3a2a* on chromosome 7. B) Mutant animals were genotyped for polymorphisms at 31609203 (A_z_), 29087847 (B_z_), 28090051 (C_z_), 26124850 (D_z_), and 19034592 (E_z_) bp to identify recombinants and predict the distance to the causative mutation. Points representing marker locations, and horizontal lines representing the estimated distance to mutation are overlaid on the SNP index for the interval. C) Representative images of larvae after editing at *slc3a2a* and *slc3a2b* loci; non-injected larvae, as well as *zion*(+/?) and *zion*(-/-) siblings are shown for comparison (dark yolk 17±9%, N = 9, n. D) *slc3a2a*^c1001/+^ in-cross generates larvae with the dark yolk phenotype; dark yolk frequency is shown as mean ± standard deviation, N= 3, n= 876. E) *slc3a2a*^c1001/+^ crossed to *zion*(+/-) generates larvae with the dark yolk phenotype; dark yolk frequency is shown as mean ± standard deviation, N = 3, n = 299. For panels D-F, animals are 5 dpf, scale bar represents 1 mm. F) ApoBb.1-nanoluciferase quantification in *zion* mutants and siblings. Mean +/- standard deviation, N = 3, n = 2-14, outliers were removed by the ROUT method (Q = 1%). P<0.05 by two-way ANOVA with Geisser-Greenhouse correction and Tukey’s multiple comparisons test. * vs. +/- and +/+. G) HuGE scores for SLC3 and SLC7 genes quantify the association of human variants with serum triglyceride (Tg) and total cholesterol (Chol). Genes with established links to B-lp synthesis are shown for comparison. HuGE association categories are noted on the right.

**Table 2:**
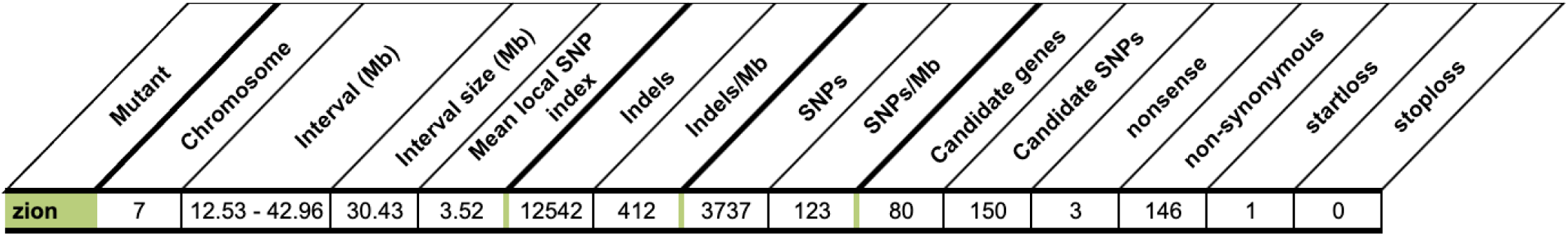
*zion* WGS summary.

To further confirm *slc3a2a* as the causative locus, CRISPR was used to generate an *slc3a2a* mutant with a 26 bp deletion in exon 4 that removes the splice acceptor site and part of the exon (Extended Data Fig. 6D); we named this allele c1001. Incrossing *slc3a2a*^+/c1001^ fish produces larvae with the dark yolk phenotype (Fig. 5D). The frequency of dark yolk in c1001 was slightly sub-mendelian suggesting the c1001 allele may be more mild and not fully penetrant. Importantly, the c1001 allele fails to complement *zion*, as outcrossing *slc3a2a*^+/c1001^ to *zion*^+/-^ also produces dark yolk larvae in the predicted mendelian ratio (23±5%) (Fig. 5E). The same result was observed for two additional alleles of *slc3a2a* (c1002, c1003) (Extended Data Fig. 6E-F). The C>T mutation in *zion* mutants was confirmed by Sanger Sequencing of genomic DNA (Extended Data Fig. 6G) and a genotyping protocol was developed. *slc3a2a*^zion^ larvae had significantly fewer B-lps after phenotype onset (4 dpf) relative to *slc3a2a*^+/*zion*^ and *slc3a2a*^+/+^ siblings (Fig. 5F). The human ortholog of *slc3a2a*, SLC3A2, heterodimerizes with SLC7 family members to form amino acid exchangers^35^. To further evaluate the potential role for SLC3A2 in B-lp metabolism we evaluated SLC3A2 and SLC7 genes for polymorphisms with associations with abnormal lipid metabolism parameters using the HUGE score calculator^36^. While SLC3A2 itself is not associated with dyslipidemic phenotypes, several of its binding partners (SLC7A2, SLC7A6, SLC7A10, and SLC7A13) are (Fig. 5G). Taken together, these data illustrate the power and efficiency of the WheresWalker pipeline for mutation mapping.

## DISCUSSION

Here we introduce WheresWalker, a mutation mapping tool composed of three parts: 1) a mapping-by-sequencing algorithm that identifies a genomic interval containing the causative mutation, 2) a recombinant mapping module that utilizes background polymorphisms to refine the bioinformatically defined interval to narrow the list of candidate genes, which can be 3) tested using high efficiency CRISPR/Cas9 reverse genetics (Fig. 6). This hybrid model leverages the best of both traditional and contemporary approaches to enable rapid mutation mapping on the order of weeks, as opposed to years. We rigorously test the mapping-by-sequencing component of WheresWalker by identifying 3 loci from a recent mutagenesis screen in zebrafish, and show that WheresWalker can also be applied to map mutations in maize, and likely many other species. Consistent with modeling of sequencing coverage in Arabidopsis with the SHOREmap tool^37^, we find the number of candidate mutations is reduced with increased coverage. Based on our data, we recommend sequencing at ∼30X coverage to identify a high-quality interval with the fewest SNP candidates. Our data strongly supports the use of multiple parental pairs when generating DNA pools for sequencing. While not essential, using outcrossed parents did improve the ratio of signal to noise.

**Figure 6:**
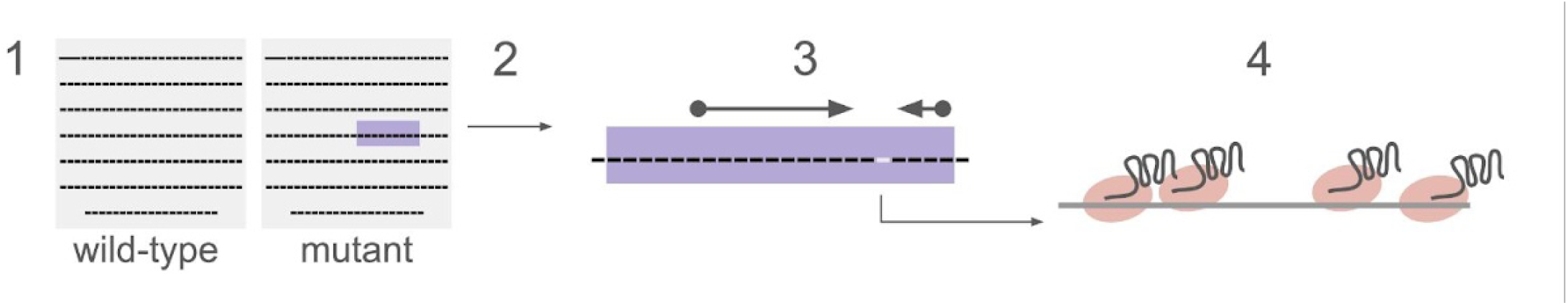
WheresWalker Pipeline. WheresWalker utilizes whole genome sequencing from wild type and mutant pooled genomic DNA (1) to identify an interval of homozygosity in the mutant genome which contains the causative mutation (2). Indel locations are outputted for use in recombinant mapping to narrow the interval (3). Gene candidates are tested using a redundant CRISPR approach (4).

Using WheresWalker, we map a novel dark yolk mutant, *zion*, to a nonsense mutation in *slc3a2a* and show that this mutation leads to reduced levels of B-lp suggesting a defect in B-lp biogenesis. To our knowledge, there is currently no published literature linking *slc3a2a* directly to B-lp metabolism. SLC3A2 itself was not associated with dyslipidemia, but several SLC7 dimer partners were, suggesting the link between *slc3a2a* function and B-lp metabolism may be specific to a subset of transported substrates (e.g. amino acids), tissue expression patterns, and/or subcellular localization. Amino acid availability may play a direct role in B-lp synthesis by providing substrate for the synthesis of ApoB (a 4563 aa protein in humans and a 3730 aa protein in zebrafish). Many amino acids (Ala, Arg, Asn, Gln, His, Leu, Met, Ser, Thr, and Val) activate mTORC1 activity^38^. mTORC1 activity is associated with increased secretion of B-lps^39,40^ and has been linked to metabolic disease states^41,42^. Specifically, the SLC3A2/SLC7A5 leucine transporter has been shown to be important for mTORC1 activation^43^. Further characterization will be required to better understand the mechanistic role of SLC3A2 in B-lp metabolism. These data demonstrate the power of unbiased forward genetic approaches to assign new functions to genes.

CRISPR has emerged as a powerful tool for reverse genetics, but has increasingly been used for genome-wide screening to generate null alleles (CRISPRko)^17,44–46^. However, forward genetic ENU/EMS mutagenesis screens remain relevant for their propensity to generate null as well as hyper- and hypo-morphic alleles, a functionality that is not yet available for high-throughput CRISPR-based technology. Moreover, chemical mutagenesis can be deployed in organisms for which CRISPR or other mutagenesis strategies have not been developed. Historically, the major drawback of chemical mutagenesis approaches has been the years required to identify the gene/causative mutation(s) underlying exciting new phenotypes. Existing mapping-by-sequencing tools can narrow the relevant genomic region, but still may select an intractable number of candidate genes. Additional expertise in bioinformatics is required to extract indels if mapping is required. WheresWalker solves this problem. It is an efficient, easy-to-use, and freely available mapping and positional cloning pipeline that requires virtually no bioinformatics experience. Because WheresWalker utilizes background polymorphisms for interval picking and fine mapping, it is best deployed in organisms bred in polygenic backgrounds. In the current wave of emerging model organisms, we anticipate WheresWalker will be a critical tool for foundational mutagenesis screens in novel model species and can also be applied to yet unmapped mutants and modifiers from historical and contemporary screens in traditional genetic models.

## METHODS

### Fish Lines and husbandry

Adult zebrafish (*Danio rerio*) were maintained at 27°C on a 14:10 h light:dark cycle and fed once daily with ∼3.5% body weight Gemma Micro 300 (Skretting USA). Embryos were obtained by natural spawning and were raised in embryo medium at 28.5°C and kept on a 14:10 h light:dark cycle. On 3 dpf, plates were cleaned and fresh embryo medium was added. All lines including *slc45a2*^b4^, Fus(ApoBb.1-nanoluciferase) (Fus(ApoBb.1-nluc)), *mttp*^c655^, *mttp*^stl^ , *apobb.1^wz25^, mia2^mw91^*, and *dgat2*^sa13945^, were maintained in the AB background. Selected lines were outcrossed to WIK for mapping. All Zebrafish protocols were approved by the Carnegie Institution Department of Embryology Animal Care and Use Committee (Protocol #139).

### Brightfield Imaging

3-5 dpf larvae were immobilized in cold 3% methylcellulose and imaged using a Nikon SMZ1500 microscope with HR Plan Apo 1x WD 54 objective, Infinity 3 Lumenera camera and Infinity Analyze 6.5 software.

### ENU mutagenesis

ENU mutagenesis was performed as previously described^47^ with some modifications. Specifically, 20 healthy and fecund AB male zebrafish were exposed to ENU (Millipore-Sigma, N3385) prepared at 3.3 mM in 10mM sodium phosphate buffer pH 6.5 in system water for 1 hour in a dark, quiet room. Tricaine (Millipore-Sigma, A5040) was added to a final concentration of 15 mg/L for the final 5 minutes of mutagenesis to minimize stress during mutagen removal. Mutagen was removed by draining the contaminated solution to a minimal volume (∼100 mL) and diluting with wash solution (1.9L, 10 mg/mL tricaine in 10 mM phosphate buffered system water). After 3 washes, exposed fish were allowed to rest for 2 hours in wash solution before being transferred in a minimal volume (∼100 mL) to an on-system 10 L tank with constant system water inflow and off-system drainage. After 48 and 96 hours, mutagen exposure was repeated for a total of 3 exposures. Mutagenized fish were rested 2 weeks before being bred to clear mutagenized sperm. All males survived initial ENU exposure, 17 survived past the 2 week resting period, and 13/17 remained fecund post-mutagenesis. ENU mutagenesis was performed in a dedicated space with appropriate personal protective equipment. All ENU contaminated solutions and materials were neutralized in a deactivation solution of 10% sodium thiosulfate 1% sodium hydroxide.

### Propagation and Screening

Mutagenized (F0) males were crossed over two weeks to clear mosaic post-mitotic germ cells^23^ before outcrossing to *slc45a2*^b4^ to test for mutational load or to AB females to generate F1 fish. F1 fish were outcrossed to Fus(ApoBb.1-nluc)^+/-^ to generate F2 families. On 5 or 6 dpf, up to 50 larvae were put on the system to be raised for each F1. Once mature, F2 siblings were incrossed to generate F3 clutches for screening. Up to 100 larvae/clutch were collected and observed under a stereoscope using transmitted light at 3 and 5 dpf. On 5 dpf, larvae were anesthetized with Tricaine solution so that the yolks could be more easily assessed. An F2 family was considered fully screened once 6 unique clutches had been observed. F2 families that produced dark yolks at ∼25% in one or more clutches were considered mutant families and were named and prioritized for characterization.

### Whole genome sequencing

Larvae were pooled and flash frozen on dry ice then stored at -20°C. Genomic DNA was extracted using the DNeasy Blood & Tissue Kit (QIAGEN, 69504) according to the included protocol for Tissue. To achieve sufficient lysis, reagents were scaled 1.5X for pre-wash steps and the proteinase K digest was 15 minutes. DNA was eluted in nuclease-free water. DNA was prepared for sequencing using the Illumina DNA Prep, Tagmentation kit (Cat#20018705) and Nextera DNA CD Indexes (Cat#20018707) using an input of 200 ng and 5 PCR cycles per the user manual. Sequencing was performed on an Illumina NextSeq500 as a 150 bp single-end run with dual (8x8 bp indexing). ∼30X coverage (∼268 million reads) was acquired for each sample. The POLCA genome polishing software from the MaSuRCA genome assembly and analysis toolkit was used to align reads to the GRCz11 genome assembly and identify variance. Fastq files for published maize datasets were downloaded from the SRA database (SRR7467444, SRR7467445) and processed by POLCA as described above.

### WheresWalker

The WheresWalker pipeline is implemented in Perl and bash scripting languages. The pipeline is designed to identify a region of relative homozygosity in the mutant population, compared to the wild type population. The pipeline then analyzes the variants in the region using a modified version of the ANNOVAR software with the annotation and the transcript sequences as input. The key technique used in the pipeline is that instead of looking for a homozygous region, it looks for a region that is *less heterozygous*, utilizing the SNP data for the mutant and wild type. The data for mutant and wild type is input in the form of VCF (Variant Call Format) files. The pipeline then computes a value of SNP index (see Results) for every 10000 bp window in the genome. The SNP index is then smoothed by a moving average function with the window size of about 1 cM (750 kbp), or 75 10000 kb windows. The pipeline looks for a global maximum of the smoothed SNP index over the entire genome sequence. The beginning and the end of the single window are then identified by looking for SNP index values below 0 around the maximum. The pipeline then analyzes all mutations inside the window with the ANNOVAR software and annotates mutations with their consequences. Mutations are reported based on allele observation filtering parameters, where RO is the number of Reference observations (the number of reads that agree with the genome sequence) and AO is the number of Alternative observations (the number of reads that disagree with the genome sequence at that locus). A mutation is reported if the mutant RO < 2 and AO ≥ 2 and one of the following: 1) the mutation is heterozygous in the wild type sample (1> RO/AO < 4), 2) the mutation is present only in the mutant dataset. The most attention should be paid to non-synonymous/frameshift/start loss/stop loss mutations. Table 1 lists the number of such mutations in Experiments described in this study. Optionally WheresWalker can add SIFT values to each mutation, which helps to exclude well-studied cases. WheresWalker output files were analyzed and plotted in R (supplementary file 1).

### PCR

Genomic DNA for PCR was prepared by incubating whole larvae or adult fin clips in 50-100 µL 50 mM NaOH at 95°C for 15-20 min followed by neutralization with a 10% volume of 1 M Tris pH 8.0. Primers were designed using NCBI Primer Blast. PCR reactions were prepared using 4 µL Green GoTaq® Flexi Buffer, 1.5 µL 25 mM MgCl_2_, 0.1 µL GoTaq® Flexi DNA Polymerase (Promega, M8295), 0.5 µL dNTP Mix (Qiagen, 201901), 0.5 µL each of forward and reverse primers (10 µM, synthesized by Eurofins Scientific), and 1 µL of genomic DNA with nuclease-free water to 20 µL. PCR was performed for 35 cycles. Assay specific annealing temperatures, extension times, and oligo sequences are reported in supplementary file 2. PCR products were separated on a 2-3% agarose (Millipore Sigma, 11685678001) gel containing 0.5 µg/mL ethidium bromide (ThermoFisher, 15585001) and visualized with UV excitation.

Genotyping primers were designed using dCAPS finder 2.0^48^ to introduce a restriction enzyme cut site into either the wild type or mutant PCR product. The *arches* allele was genotyped using primers SFMRF111 and SFMRF110^dCAPS^, the 205 bp product was digested with ApoI (NEB, R0566) and incubated for 1 h at 50°C to generate 205 bp (wild-type) and 179 bp (mutant) products. The *zion* allele was genotyped using primers SFMRF344^dCAPS^ and SFMRF346, the 116 bp product was digested with BccI (NEB, R0704) for 1 h at 37°C to generate 83 bp (wild-type) and 116 bp (mutant) products. Mttp alleles^21^ and ApoBb.1-nluc^24^ were genotyped as previously described. All primer/oligo sequences are reported in supplementary file 2.

### Recombination calculations

Recombination frequency (Rf) was calculated as the fraction of animals with recombination out of the total number of animals observed multiplied by 100. Rf*cM was used to estimate the distance to the causative locus; a cM is 0.74 Mb in zebrafish^15^. In some cases, for the *zion* mutants, genotype could not be determined by gel because the primers failed to adequately amplify the region, likely due to inefficient primer binding or amplification (Extended Data Fig. 5A-B). In some cases, unknown genotypes could be inferred based on the genotype of adjacent markers. Inferred genotypes were considered when calculating the average estimated position of the causative locus and are reported in the main text. Estimated positions calculated with only empirical genotypes are reported in Extended Data Fig. 5C.

### Sanger Sequencing

PCR amplicons were prepared by standard PCR then sent to Genewiz for sanger sequencing. The *arches* amplicon was prepared with primers SFMRF111 and SFMRF104, and sequenced with SFMRF111. The *zion* amplicon was prepared with primers SFMRF460 and SFMRF346 and sequenced with SFMRF460. The *slc3a2a*(c1001) amplicon was prepared with primers SFMRF827 and SFMRF349 and sequenced with SFMRF347. All oligo sequences are reported in supplementary file 2.

### Lipoglo Assays

ApoBb.1-nluc quantification and visualization was performed as previously described^24^. Briefly, for ApoBb.1-nluc quantification, animals were homogenized by sonication in a microplate-horn sonicator (Qsonica, Q700 sonicator with 431MPX microplate-horn assembly) in 100 µL stabilization buffer (1 g sucrose, 400 µL 0.5 M EGTA pH 8.0, 1 cOmplete™, Mini, EDTA-free Protease Inhibitor Cocktail (Millipore Sigma, 11836170001), in 10 mL deionized water). 4-40 µL homogenate was diluted in reaction Buffer (0.5% Nano-Glo® Luciferase Assay substrate, 25% Nano-Glo® Luciferase Assay Buffer in PBS) to a final volume of 80 µL in an OptiPlate™-96F Black flat-bottom plate. A SpectraMax M5 (Molecular Devices) or Tecan Spark plate reader set to top read with 500 ms (SpectraMax) or 20 ms (Tecan Spark) integration time was used to collect luminescence measurements. For ApoBb.1-nluc visualization in whole-mount animals, embryos were fixed in 4% paraformaldehyde (Electron Microscopy Science, 15714, diluted in PBS) for 3 h at room temperature then washed 3x15 minutes in 0.1% tween-20 (ThermoFisher Scientific, J66278.AP, diluted in PBS). Embryos were immobilized in 1% low melt agarose (Fisher Scientific, BP160, prepared in 1x TBE) containing 1% Nano-Glo® Luciferase Assay substrate. Images were collected on a Zeiss Axiozoom V16 microscope V16 with a Zeiss AxioCam MRm. Luminescent signal was collected over 30 seconds with no illumination.

### Guide synthesis and F0 CRISPR

Guide template oligo sequences were obtained from Wu et. al.^17^; template oligo and CRISPR tail primer sequences are reported in supplementary file 2. Oligos were synthesized by Eurofins Scientific. Guide RNA was prepared according to Wu et. al.^17^ with some modifications. Template DNA was prepared by PCR with CRISPR tail primer and Phusion™ High Fidelity Polymerase (ThermoFisher Scientific, F530). Template DNA was pooled according to target (*slc3a2a* = SFMRF324-327, *slc3a2b* = SFMRF404-407) then cleaned using QIAquick PCR Purification Kit (Qiagen, 28104). MEGAshortscript™ T7 Transcription Kit (ThermoFisher, AM1354) and 200-1000 ng template DNA was used to generate gRNA. gRNA was isolated by alcohol precipitation according to the kit protocol. The gRNA pellet was reconstituted in nuclease-free water, diluted to 2500 ng/µL, aliquoted, and stored at -80°C. An injection mix of 300 mM KCl, 0.1% phenol red, 0.8 µg/µL Alt-R™ S.p. Cas9 Nuclease V3 (IDT, 1081058) and 1000 ng/µL pooled gRNA in nuclease-free water was prepared. 2 nL of injection mix was injected into 1-cell stage zebrafish embryos. Larvae were raised in EM and observed on 3 and 5 dpf for yolk phenotypes.

### *slc3a2a* line generation

Guides targeting *slc3a2a* sites 2 and 3 (SFMRF325, SFMRF326) were synthesized and injected as described above except that guides were not pooled during synthesis and were combined at 500 ng/µL each in the final injection mix. Injected, F0 fish were raised to adulthood then in-crossed so that progeny could be screened for presence of the dark yolk phenotype. F0 pairs that produced dark yolk were outcrossed to AB and F1s were raised to adulthood. Genomic DNA was obtained from adult F1 fin clips to identify animals with editing at the *slc3a2a* locus by PCR amplifying a region that included guide cut sites 2 and 3 using primers SFMRF347 and SFMRF349. A male and female F1 with ∼30 bp deletions were selected for further characterization. F1 fish were incrossed, generating F2 larvae with dark yolk. F1 and mutant F2 DNA was submitted for Sanger sequencing. F2 larvae were homozygous for a 26 bp deletion which removes part of exon 4, including the splice site acceptor. The same allele was detected in F1 parents which was detected using Poly Peak Parser^49^. We named this allele *slc3a2a*^c1001^. F1 *slc3a2a*^c1001/+^ were crossed to *zion*^+/-^ to test for complementation. The same approach was used to generate, identify, and characterize alleles c1002 and c1003.

### Additional Software and Databases

Google Sheets/Docs/Slides, Paperpile, SnapGene, Prism, and R were used to prepare materials for publication. Essential public databases include NCBI^50^, Ensembl^51^ and ZFIN^52^.

## DATA AVAILABILITY

WGS datasets generated will be made publicly available from the NCBI Sequence Read Archive (SRA) upon publication. Accession numbers will be provided in supplementary file 3.

## CODE AVAILABILITY

WheresWalker is publicly available on github at https://github.com/alekseyzimin/WheresWalker.

## Supporting information

R code for analysis of WheresWalker outputs

Oligo sequences and PCR parameters

## ACKNOWLEDGEMENTS

This work was supported by grants from the National Institutes of Health: F32 GM144223 (M.F.), R01 DK093399 (S.A.F), R01 GM63904 (S.A.F), and R01 HL158054 (S.A.F). Additional support for this work was provided by the Carnegie Institution for Science endowment and the G. Harold and Leila Y. Mathers Charitable Foundation (S.A.F). The authors acknowledge Dr. Rebecca Burdine for providing WIK zebrafish and the Carnegie Embryology Sequencing Core facility, particularly Allison Pinder and Frederick Tan, for supporting sequencing efforts. In addition, the authors acknowledge Jasmine James, Tye Chicha, Victoria Murphy, and Camille Coffey for phenotyping screen mutants during lab rotations, and Julia Baer who managed the fish facility during the screen.

## AUTHOR INFORMATION

### Contributions

WheresWalker was conceived by AVZ, SLS, MF, and SAF; AVZ constructed all code. The dark yolk screen was conceived by SAF, MHW, and MF. MF, SA, NP, MK, YS, VSG, VPT, HK, and NVL, conducted all zebrafish husbandry to support the screen. MF, MRH, JLA, MS, and MHW, screened all mutant families. Characterization and analysis of *arches* was conducted by SA and MF. Characterization and analysis of *zion* conducted by NP and MF. MF, MRH, and MHW, conducted characterization for other mutants. MF and AVZ performed all other data analyses. MF wrote the original draft of the paper with review and editing by AVZ, SLS, MHW and SAF.

## ETHICS DECLARATION

The authors declare no conflicts of interest.

## EXTENDED DATA

See extended data figures 1-6.

## SUPPLEMENTARY DATA

Supplementary file 1 - R analysis file

Supplementary file 2 - oligo sequences

Supplementary file 3 - to be prepared upon SRA data submission

---

**Extended Data Figure 1:**
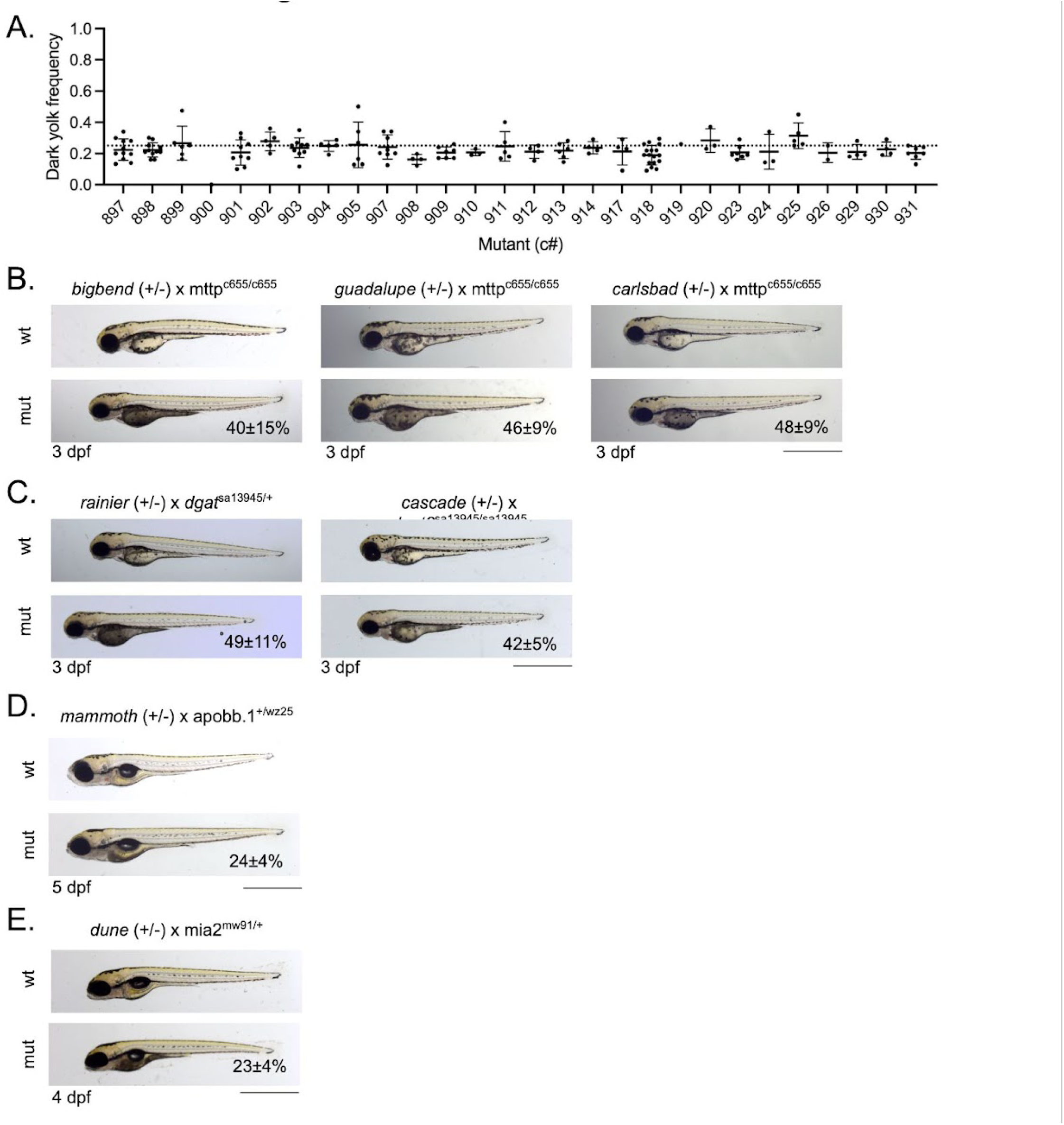
A) Phenotype frequency for all mutants identified, N = 1-17 clutches/mutant. Bars represent mean ± standard deviation. Dashed line indicates the expected frequency of 0.25. B-E) Representative images for 7 additional mutants that fail to complement known dark yolk loci, including 3 *mttp* (B), 2 *dgat2* (C), 1 *mia2* (D), and 1 *apobb.1* (E) alleles. Representative wild-type (wt) and mutant (mut) yolk phenotypes are shown. Animal age is noted. Phenotype frequency is reported as mean ± standard deviation. *bigbend*: N = 5 clutches, n = 422 animals; *guadalupe*: N = 6 clutches, n = 598 animals; *carlsbad*: N = 8 clutches, n = 784 animals; *rainier*: N = 3 clutches, n = 210 animals; *cascade*: N = 4 clutches, n = 402 animals; *mammoth*: N = 3 clutches, n = 282 animals; *dune*: N = 4 clutches, n = 180 animals;. Scale bar represents 1 mm.

**Extended Data Figure 2:**
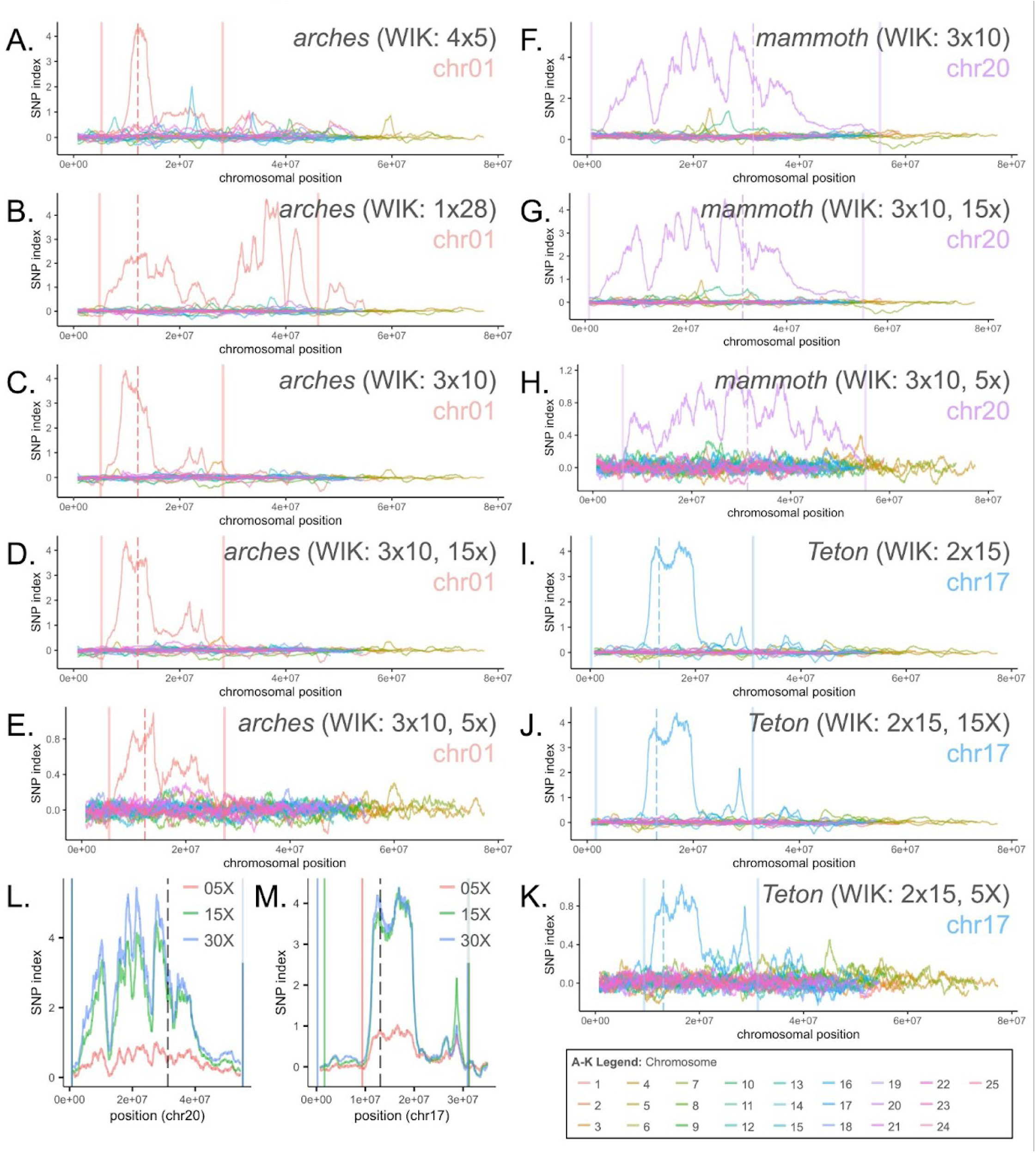
A-K) Profile of SNP index across all chromosomes corresponding to regional plots in Figure 3. Solid lines indicate left and right bounds of the interval selected by WheresWalker, dashed lines indicate the position of the causative locus. L-M) regional SNP index profile for datasets at 05X, 15X, or 30X genomic coverage for *olympic* (L) and *teton* (M) mutants. Solid lines indicate left and right bounds of the interval selected by WheresWalker, dashed lines indicate the position of the causative locus.

**Extended Data Figure 3:**
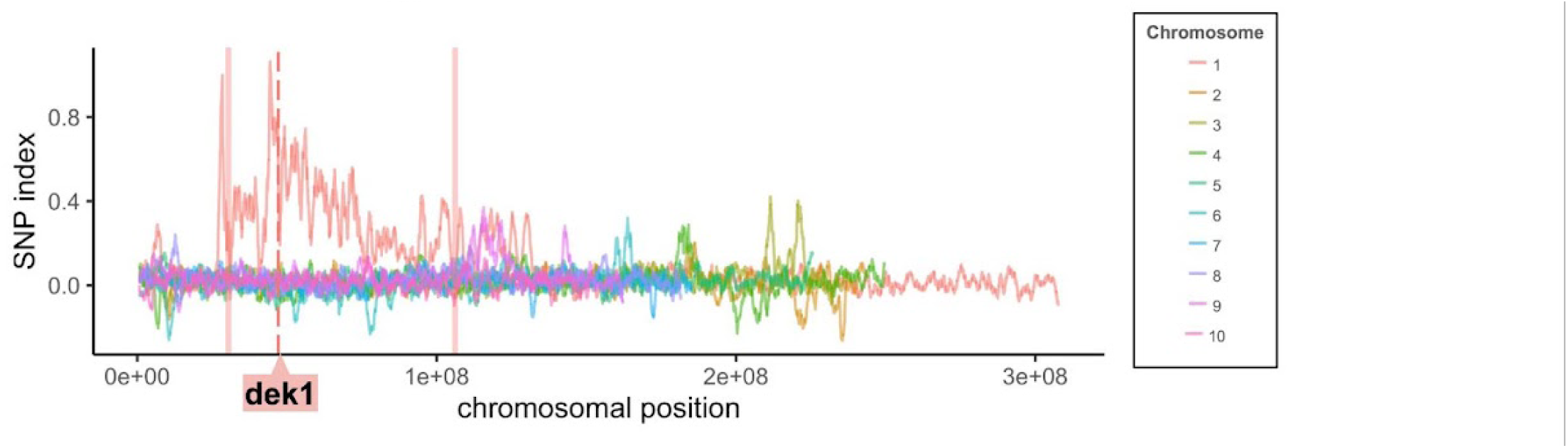
A) WGS data from vsn maize mutants and siblings were submitted to WheresWalker. The SNP index across all chromosomes is plotted. WheresWalker identifies an interval containing the causative gene, *dek1* (LOC542509). Solid lines indicate left and right bounds of the interval selected by WheresWalker, dashed line indicates the position of the causative locus.

**Extended Data Figure 4:**
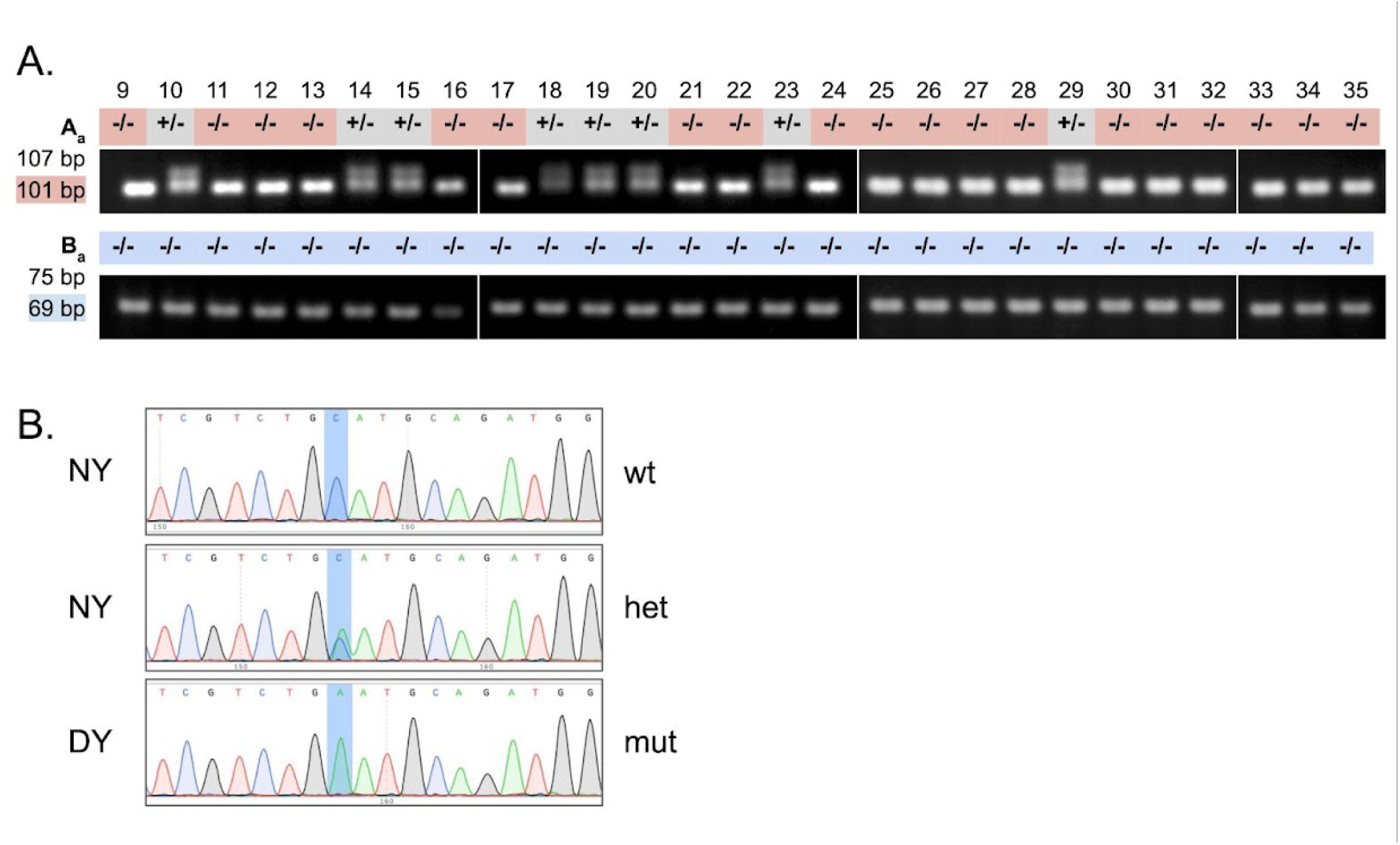
A) genotyping gels for markers A_a_ and B_a_ for *arches* mutants 9-35. PCR product sizes for wild type and mutant (highlighted) products are as indicated. B) Sanger Sequencing of normal (NY) and dark yolk (DY) animals from an *arches*(+/-) in-cross have the expected genotypes at the mutant position: wild type (wt) - C, heterozygous (het) - C/A, mutant (mut) - A.

**Extended Data Figure 5:**
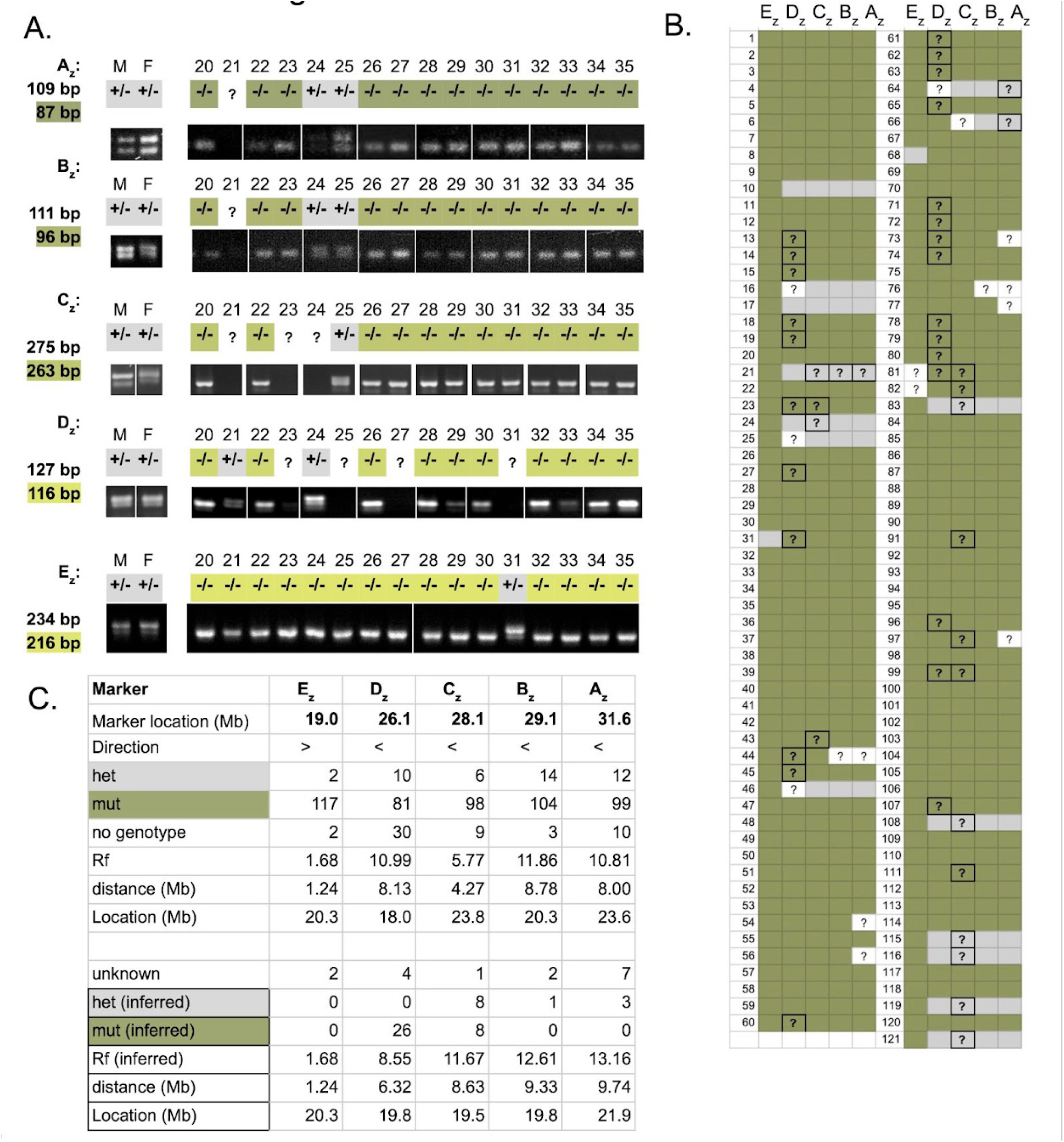
A) recombination gel for *zion* parents (M: male, F: Female) and selected *zion* mutant progeny at markers A_z_-E_z_. Genotype score is as indicated above each lane. PCR product sizes for wild type and mutant (highlighted) products are shown. B) Marker genotype score for all *zion* mutant progeny. Green shading indicates mutant genotype, gray shading indicates heterozygous genotype, white background indicates genotype could not be determined. Outlined boxes with “?” indicate the genotype could not be determined by gel, but could be inferred based on the genotype of the surrounding markers for that animal. C) Summary of genotype at each marker with calculations for recombination frequency (Rf) and estimated distance to mutation calculated with and without inferred genotypes.

**Extended Data Figure 6:**
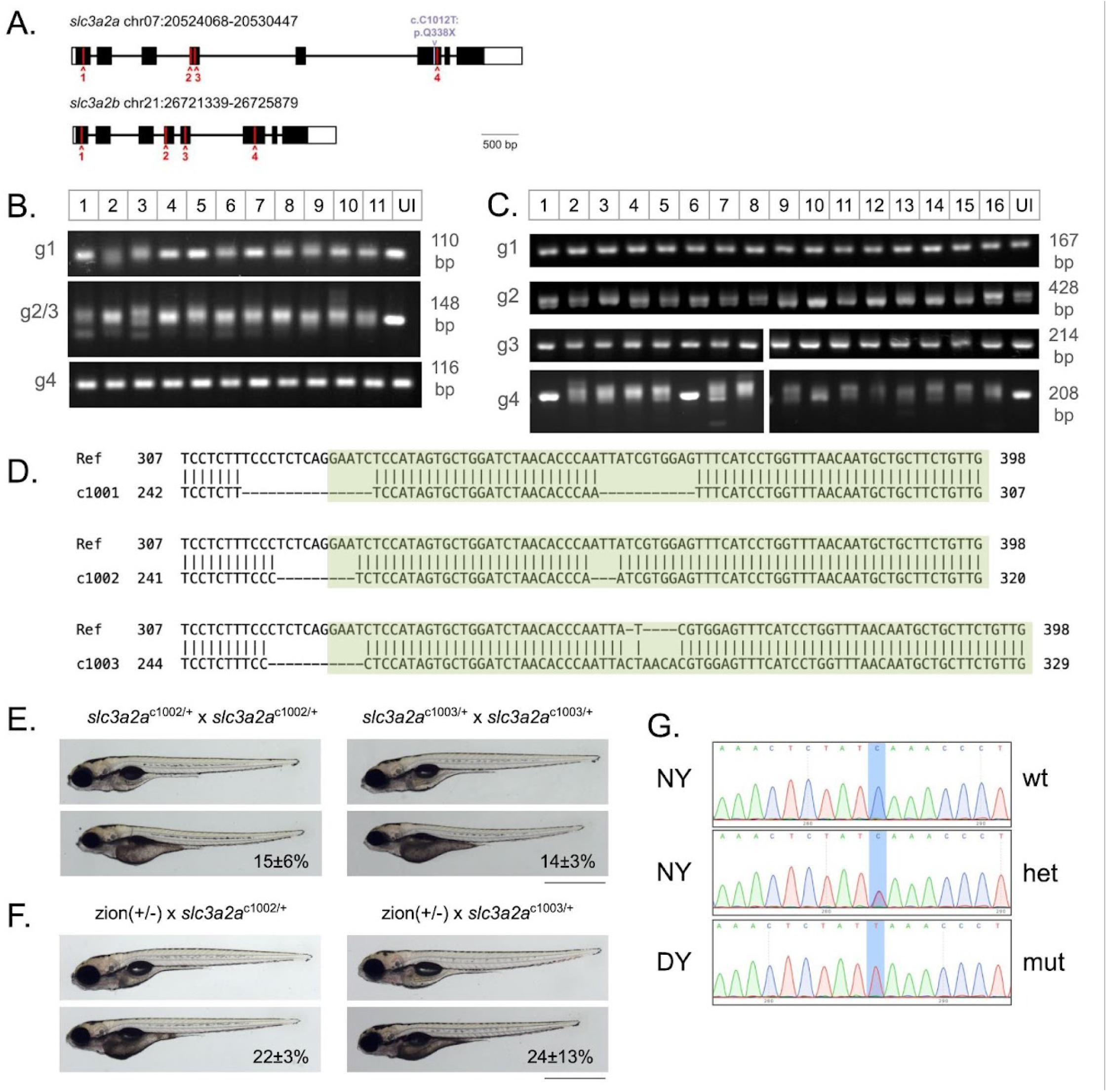
A) Schematic of the *slc3a2a* and *slc3a2b* genes. The location of the C>T base pair change in *slc3a2a* in the *zion* mutants is shown in purple. Red carats indicate locations targeted by CRISPR guides. B-C) PCR amplification around CRISPR guide sites in *slc3a2a* (B) and *slc3a2b* (C) injected animals. An uninjected animal (UI) was genotyped for comparison. “g#” indicates the guide site that is amplified for the respective gene, the expected size for each product is indicated on the right of each gel. For *slc3a2a*, editing is observed in many animals at g1 and g2/3; for *slc3a2b*, editing is observed in many animals at g4. D) c1001 (-26 bp), c1002 (-12 bp), and c1003 (-11+5 bp) *slc3a2a* alleles exhibit insertions/deletions in exon 4 and the exon 4 splice site as determined by Sanger Sequencing; exon 4 is highlighted in green. E) In-crossing *slc3a2a*^c1002/+^ (N = 2, n = 558) or *slc3a2a*^c1003/+^ (N = 2, n = 387) CRISPR mutants produces offspring with dark yolk. F) Outcrossing *zion*(+/-) to *slc3a2a*^c1002/+^ (N = 3, n = 299) or *slc3a2a*^c1003/+^ (N = 3, n = 219) produces offspring with dark yolk. For E-F, animals are 5 dpf, dark yolk frequency is reported as mean ± standard deviation, scale bar represents 1 mm. G) Sanger Sequencing of normal (NY) and dark yolk (DY) animals from a *zion*(+/-) in-cross have the expected genotype at the mutant position: wild type (wt) - C, heterozygous (het) - C/T, mutant (mut) - T.

## Notes

### Competing Interest Statement

The authors have declared no competing interest.

